# Reprogramming Epiblast Stem Cells into Pre-Implantation Blastocyst Cell-like Cells

**DOI:** 10.1101/2020.09.29.318279

**Authors:** Kiichiro Tomoda, Haiming Hu, Yoshiki Sahara, Hashimita Sanyal, Minoru Takasato, Cody Kime

## Abstract

Recently, a new wave of synthetic embryo systems (SESs) have been established from cultured cells toward efficient and ethical embryonic development research. We recently reported our epiblast stem cell (EPISC) reprogramming SES that generates numerous blastocyst (BC)-like hemispheres (BCLH) with pluripotent and extraembryonic cell features detected microscopically. Here, we further explored the system over key time points with unprecedented single-cell RNA sequencing (scRNA-seq) analysis and revealed broad induction of the 2C-like reporter *MERVL* and RNA velocity diverging three major population regions with genetic expression resembling pluripotent epiblast (EPI), primitive endoderm (PE), and trophectoderm (TE). Enrichment of those three BC-like cell fates involved key regulons, zygotic genome activation (ZGA) related genes, specific RNA splicing, and select cells meaningfully distinguished critical regulons of model cells. This analysis confirms the induction of the extraembryonic cell populations during the reprogramming and we anticipate that our unique BCLH SES and rich data may uncover new facets of cell potency, improve developmental biology, and help biomedicine advance.

## INTRODUCTION

Researching early embryonic development was the basis for developmental biology and subsequent stem cell biology. In recent decades embryology have shed light on the mammalian embryo with animal models for broad significance and to address an ethically bound human embryology (Hyun et al., 2020; Rossant and Tam, 2009, 2017). At first, early embryos were used to derive various pluripotent and multipotent stem cell lines cultures with characteristics and chimeric potential analogous to the assumed origin (Evans and Kaufman, 1981; Martin, 1981). However, the ability to form synthetic embryos from cultured cells had been elusive until recent advances enabled *in vitro* Synthetic Embryo Systems (SESs).

SESs are part of a newly emerging field akin to organoids, but reflecting early embryology through ‘embryoids’ that are far more convincing (Harrison et al., 2017; Kime et al., 2018, 2019; Rivron et al., 2018; Shahbazi and Zernicka-Goetz, 2018; Zheng et al., 2019). Several SESs exist and some focus on modeling the blastocyst (BC) and it’s three layers of trophectoderm (TE), primitive endoderm (PE), and pluripotent preimplantation epiblast (EPI). Some SESs utilize embryonic stem cell (ESC) and trophoblast stem cell aggregations to model the pre/post-implantation embryo *in vitro*. Others SESs, including work from our group, involve cell reprogramming or unique cell plasticity states that give rise to BC-like cysts from single cultures. Various SES approaches building on BC-like cyst formation *in vitro* continue to be developed and explored as each system pioneers to widen embryology at large.

In a related field, reprogramming cells with exogenous factors (e.g., transcription factors, small molecules, cytokines, nutrients) pioneered new dimensions in cell biology by inducing donor cells to desirable and generally unforeseen synthetic states (Davis et al., 1987; Kime et al., 2016, 2019; Takahashi et al., 2007; Woogeng et al., 2020). Indeed, cell analogs of early embryonic BC-lineage cells have been induced (Benchetrit et al., 2019; Kubaczka et al., 2015; Parenti et al., 2016; Takahashi and Yamanaka, 2006). Our past epiblast stem cell (EPISC) reprogramming induced high-quality chimera-forming naïve-like cells with X-chromosome reactivation (Kime et al., 2016). We recently showed that the same reprogramming generated plates of BC-like hemispheres (BCLH) with KRT8+ (TROMA-I) TE-like cells surrounding the Xa/Xa EPI-like naïve ESC region which also had a PE-like GATA4+/GATA6+/PDGFRA+ population at its inner-face toward the putative blastocoel (Kime et al., 2018, 2019). However, detailed cellular gene expression and regulation of the converting BCLH cells to resemble the three BC-lineage cells was previously unknown.

The BCLH SES can be easily set up and generates BCLH efficiently from EPISC cultures. We established EPISC with the 2C-reporter *MERVL* and saw broad early expression prior to BCLH cyst-like formation. We applied single cell RNA sequencing (scRNA-seq) and saw that on Day5, three distinct regions of cells branched with gene expression resembling the blastocyst’s TE, PE, and EPI lineages. The three regions each had RNA velocity toward Day7 cells that extended those diverging regions and further enriched convincing cells of the postulated BC-cell identities. Furthermore, RNA splicing regulation and gene regulatory networks implicated significant cell reprogramming had occurred with germ and zygotic genome activation (ZGA) signature genes. Herein we detail these observations and anticipate a welcome interest in the relatively poorly explored aspect of EPISC SES reprogramming into much earlier embryonic cells.

## RESULTS

### Naïve ESC in 2iLIF may Stabilize MERVL+ Reporter Expression

For this study, we found TE and PE scRNA-seq data from previous reports, shown later, and required control naïve embryonic stem cells (ESCs). We therefore integrated BL6 ESCs with *MERVL::RFP* reporters that were cultured in our modified media on laminin as previously reported (Kime et al., 2019). Two distinct populations of ESCs stabilized; one with traditional naïve ESC dome-like morphology and transient *MERVL::RFP* expression (Macfarlan et al., 2012), and the other with a unique larger cell morphology and consistent range of *MERVL::RFP* expression (Figure 1A). Our scRNA-seq sample of the culture confirmed our suspicion of a ‘duality’ because cells clustered into two distinct groups (Figure 1B) that we bisected at the origin of UMAP_1: the left cluster termed ‘ESC’ and the right cluster with much more *MERVL::RFP* ‘2C-like’ reporter expression (Figure 1B,C) termed ‘ESC2CL’. Although technically cultured and sampled as one, the ESC2CL had higher scRNA-seq features and counts (Figure S1A). When compared, both populations generally retained similar core pluripotency features (Figure 1D) although differential gene markers could also be identified (Table S1, Table S2) with little outstanding genes each group’s top 20 (Figure S1B). For technical similarity in scRNA-seq sampling of naïve ESC controls and a common interest in ESC *MERVL* reporter activity, we used both ESC and ESC2CL clusters through this study.

**Figure 1:**
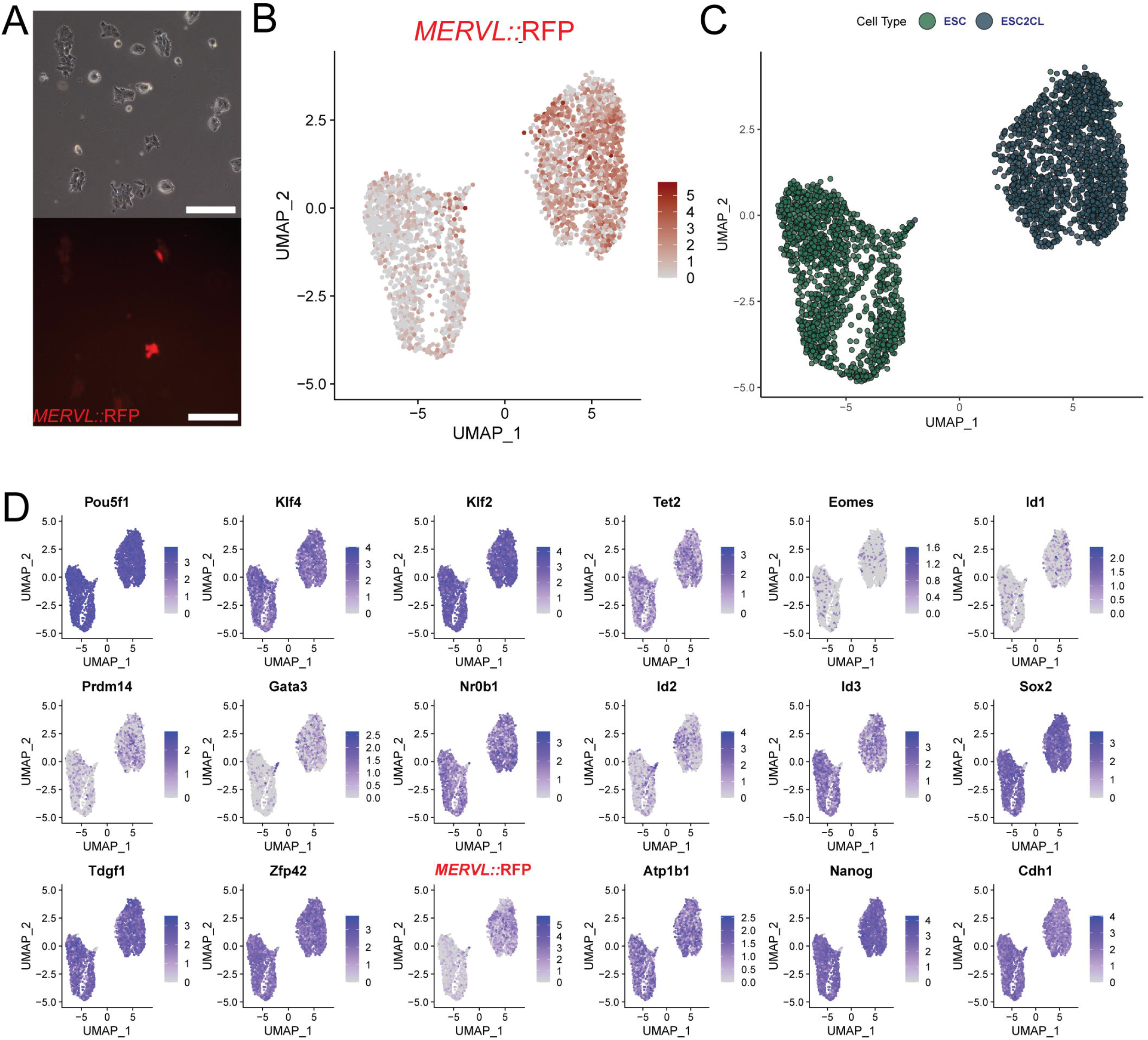
Mouse ESCs and MERVL Reporter Expression. **A)** A duality of ESC culture in 2iLIF viewed with brightfield imaging (top) and *MERVL::RFP* expression (bottom). *Scale bars = 200μm*. **B)** UMAP based gene expression feature plot for transgenic *MERVL::RFP*. **C)** UMAP plot with newly labeled ESC and ESC2CL populations. **D)** UMAP based gene expression feature plots for pluripotency genes.

### The BCLH SES Induces 2C-reporter, XGFP, and Three Regions of Blastocyst-like Lineage Cells

We previously generated (Kime et al., 2018, 2019) EPISC with XGFP and *MERVL::RFP* reporters that are completely off when viewed in fluorescence microscopy (Figure 2A). We induced BCLH reprogramming with and sampled the Day5 and Day7 reprogramming cells along with the starting EPISC and the duality ESC/ESC2CL for scRNA-seq with as standard workflow including SkewC (Abugessaisa et al., 2020) to select high quality cells for analysis (Figure S2A). On Day4 we rarely spotted XGFP while many cells showed *MERVL::RFP* activation that continued through Day5 and Day7 as XGFP activated (Figure 2B). The trend of full colonies expressing the MERVL reporter was also visible with a rapidly degrading D2nRFP (Kime et al., 2018; Li et al., 1998) (Figure S2B).

**Figure 2:**
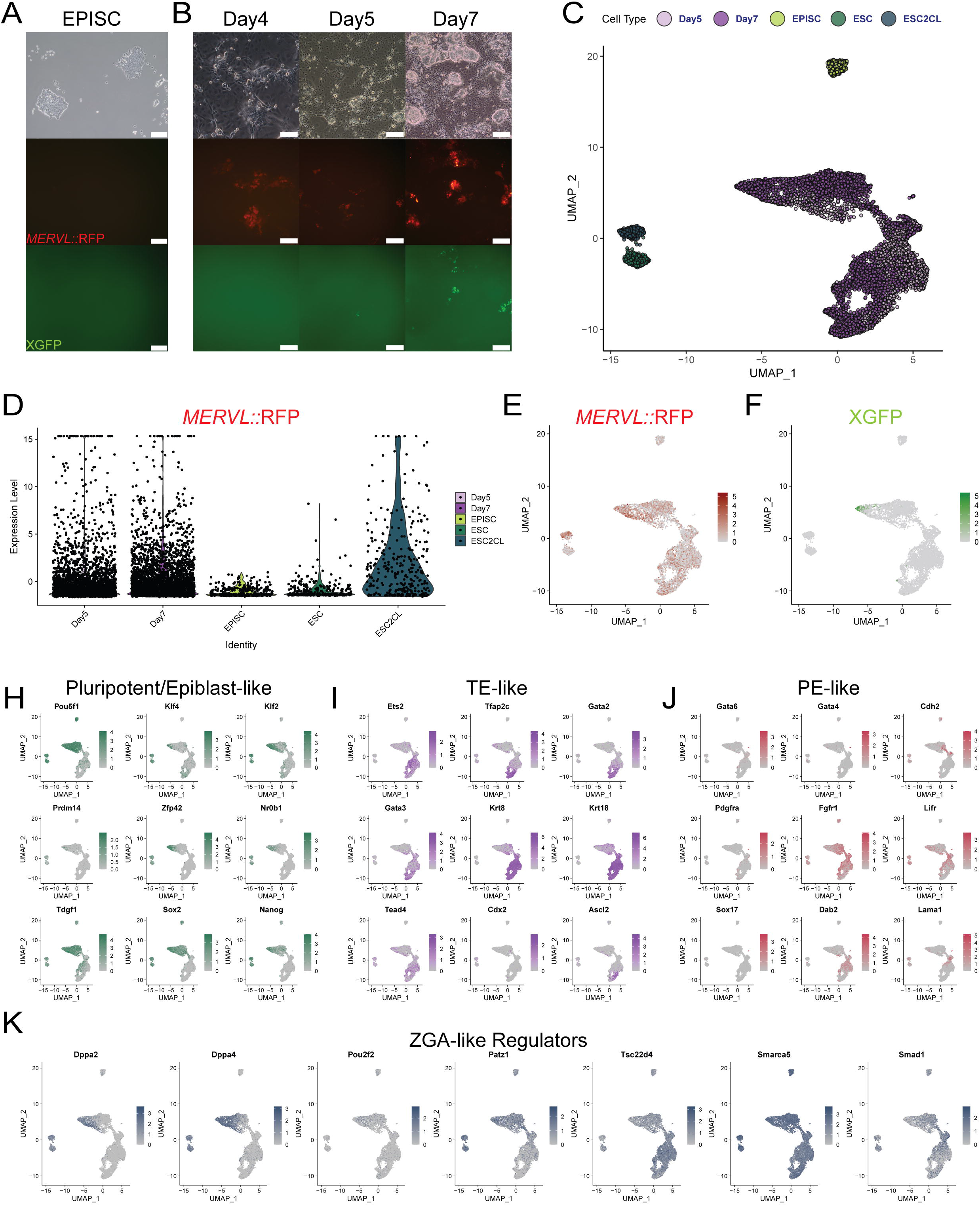
BCLH Cell Induction and Regional Gene Expression. **A)** EPISC culture with *MERVL::RFP* and XGFP reporters viewed with brightfield imaging (top) *MERVL::RFP* expression (mid), and XGFP expression (bot). *Scale bars = 200μm*. **B)** BCLH reprogramming from EPISC with *MERVL::RFP* and XGFP reporters imaged on Day4, Day5, and Day7, for brightfield (top), *MERVL::RFP* expression (mid), and XGFP expression (bot). *Scale bars = 100μm/Day4. and 200μm/Day5:Day7*. **C)** UMAP plot clustering of EPISC, ESC, ESC2CL, Day5, and Day7 samples. **D)** Violin plot of transgenic *MERVL::RFP* reporter expression in EPISC, ESC, ESC2CL, Day5, and Day7 samples. **E)** UMAP based gene expression feature plot for transgenic *MERVL::RFP*. **F)** UMAP based gene expression feature plot for transgenic XGFP. **H,I,J)** Multiple colored UMAP based gene expression feature plots for Pluripotent/Epiblast genes (H, green), TE genes (I, purple), and PE genes (J, burgundy) associated for BC-like regional likeness. **K)** UMAP based gene expression feature plots for ZGA-like regulators.

We prepared our scRNA-seq samples in Seurat (Butler et al., 2018) with SCTransform and found a clear trend of three regions of cells on Day5 that clustered closely to similar expanding and surrounding regions on Day7 (Figure 2C). In general, cells in the three regions often clustered together while Day7 cells had more distinguished separation (Figure 2C). The EPISC, the ESC, and the ESC2CL cells clustered separately (Figure 2C). Surprisingly, many reprogramming cells in all regions had enriched the *MERVL::RFP* expression (Figure 2D,E), while mostly one specific region of Day7 cells expressed the XGFP reporter (Figure 2F) grossly related to naïve pluripotency and the ICM.

Checking gene expression revealed that common late BC-lineage cell markers were enriched and relatively focused in the cells of three separated regions with Pluripotent/Epiblast-like, TE-like, and PE-like genes (Figure 2H,I,J). The Epiblast-like region colocalized with the XGFP expression (Figure 2F) and enriched for pluripotent genes (*Pou5f1, Zfp42, Tdgf1, Nr0b1, Klf2*) (Figure 2H). Importantly, XGFP was detected in the cells where Klf2, Klf4 and Prdm14 were expressed (Figure 2F,H) relating the importance of those genes to reactivate the inactive X chromosome as previously reported (Gillich et al., 2012; Kime et al., 2016). In the TE-like region, we observed numerous important TE establishing genes (*Ets2, Tfap2c, Gata2, Gata3, Elf4*) (Figure 2I) with remarkable expression of *Krt8* and *Krt18* recently reported to organize extraembryonic fate determination in the compacting and polarizing embryo (Lim et al., 2020). Also, the smaller PE-like region there had milder yet focused enrichment for important PE genes (*Pdgfra, Gata4, Gata6, Fgfr1, Lifr, Lama1*) (Figure 2J).

The 2C-like MERVL reporter in ESCs has been used in several studies yet its use in naïve ESCs has been limited due to an unclear relationship to ZGA early embryonic-like plasticity. In our SESs we found utility with this reporter (Kime et al., 2018, 2019) related to heightened cell plasticity and therefore checked numerous recently reported ZGA-like regulators and ZGA signature genes derived from powerful screens (Alda-Catalinas et al., 2020). Many ZGA-like regulators were induced in reprogramming cells (Figure 2K), and ZGA signature genes were often highly expressed broadly or regionally in Day5 and Day7 reprogramming cells (Figure S2C), while many were not expressed in the stem cell controls including the ESC2CL cells (Figure S2C). As such, MERVL reporter ESCs may have a narrower threshold to activate and reflect smaller differences while de novo MERVL reporter activation in reprogramming cells might indicate a greater composition of ZGA-like genomic remodeling.

### Three Regions of Blastocyst-likeness Enrich Over Time

To investigate the state of cells on Day5 and Day7, we employed RNA Velocity with Veloctyo (La Manno et al., 2018) to determine the ‘direction’ of cell change and view RNA splice variation. The EPISC and the ESC/ESC2CL had RNA velocities pointing inward, demonstrating stable states (Figure 3A). As we anticipated there was an obvious trend of RNA Velocity from Day5 toward Day7 among the three regions, all three of which pointed away from each other (Figure 3A). Perhaps reflecting the PE formation of a BC, the ratio of cells driving into the PE-like region was fewer and distributed between the Epiblast-like and extraembryonic-like regions (Figure 2H,J, Figure 3A).

**Figure 3:**
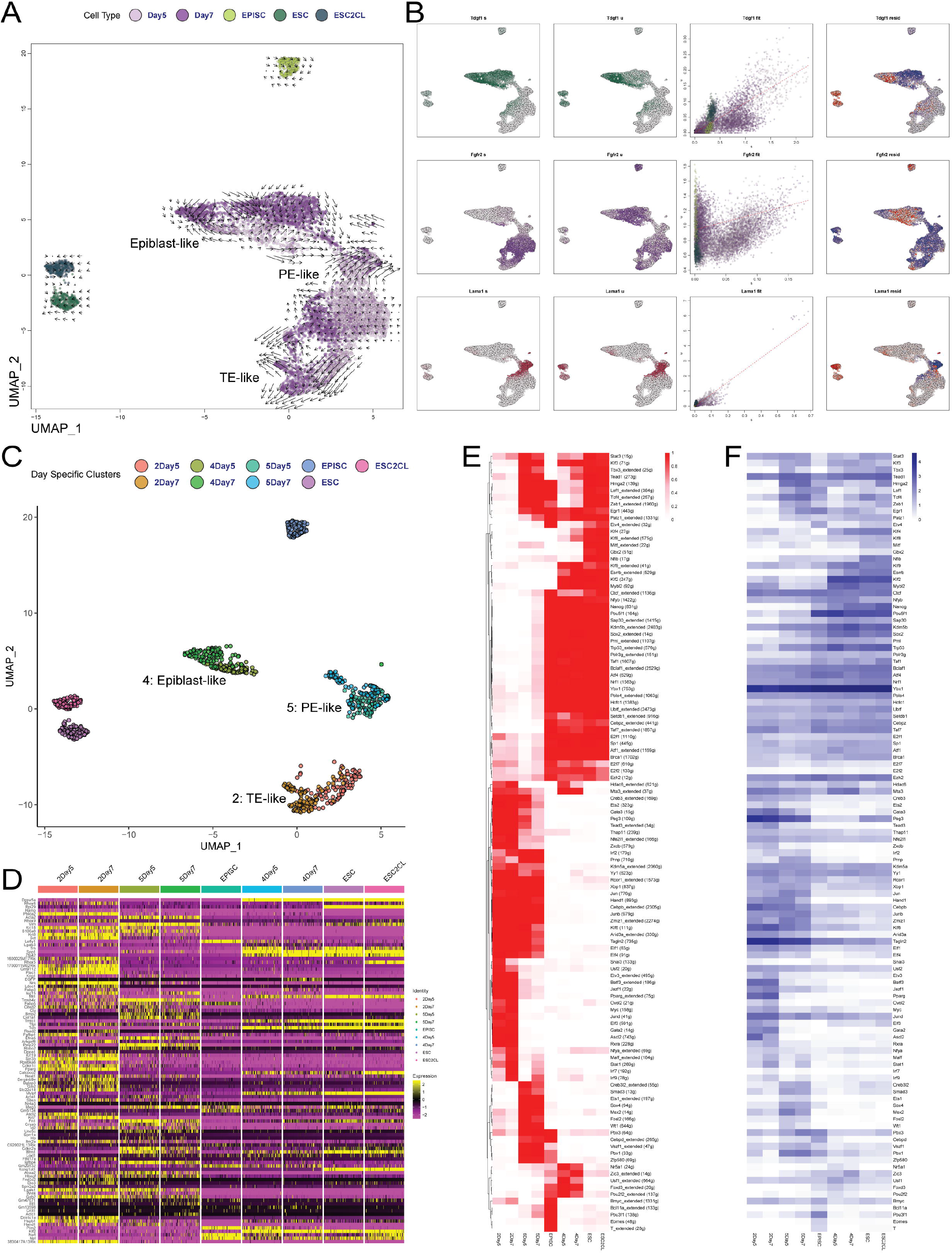
BCLH Three Region RNA Velocity and Gene Regulation. **A)** UMAP based RNA Velocity plot for EPISC, ESC, ESC2CL, Day5, and Day7 samples with BC-like regions labeled based on regional gene expression (Figure 2). **B)** UMAP based RNA splicing plots for spliced (s), unspliced (u) reads, with Cell Type coloring (Figure 3A) and residual (resid) unspliced expression shown. RNAs detected are regionally labeled by color for pluripotency (green), TE (purple), and PE (burgundy). **C)** UMAP plot of downsampled day-specific cells and control cells for use in gene heatmap (Figure 3D) and SCENIC regulon analysis (Figure 3E,F, Figure S3D). **D)** Heatmap(DoHeatmap) plot of the top 100 variable features of all cells, ordered by day-specific regions of cells and control cells. **E)** SCENIC Binarized regulons with heatmap(pheatmap) clustering and regulon activity (red scale). **F)** Heatmap(pheatmap) of the row-matched transcription factor average expression (log-transformed) for the regulons of Figure 3E.

Discriminating unspliced (u) pre-mRNA vs spliced (s) mRNAs ratios with scRNA-seq and Velocyto allows for a comprehensive understanding of important state-specific splice mechanisms beyond simple RNA detection. Tdgf1 is an early BC ICM gene (Pfister et al., 2007) often detected in ESCs, however *Tdgf1* was mostly unspliced in the ESC/ESC2CL while strongly spliced and enriched in the Epiblast-like region of reprogramming cells (Figure 3A,B) where other expression seems to implicate more ICM-like expression. The germ programming genes *Prdm1* (*Blimp1*) and *Smc1b* were also splice-enriched in reprogramming cells (Figure S3A) supporting previous consideration of the importance of germ genes toward higher plasticity in our SES reprogramming (Kime et al., 2018, 2019). As seen in base expression (Figure 2K), the ZGA-like regulators *Dppa2, Dppa4, Tsc22d4, Smarca5*, and *Smad1*, were induced across the reprogramming cells with splicing preference (Figure S3B) and the ESC2CL cells had spliced *Dppa2* more efficiently than the ESC. Similar region-specific splice mechanism preference was found with TE and PE genes *Fgfr2* and *Lama1* (Figure 3B) for translation in putative cells*. Pard6b* and *Lifr* followed a similar trend while enriched in reprogramming cells’ regions (Figure S3C). Taken together, the regulation of splice mechanisms and time-based BC-like cell region specification is apparent and related to pre-BC stage regulators.

To investigate reprogramming cells downstream gene regulation, we randomly downsampled the three regions to similar cell numbers after Seurat clusters 2:TE-like, 5:PE-like, and 4:Epiblast-like with Day-specific labeling (Figure 3C). A heatmap of the top 100 global variable genes confirmed that each cluster had a unique pattern of expression (Figure 3D). We then performed SCENIC (Aibar et al., 2017) analysis to determine cell regulons. Expectedly, each region had distinct patterning that generally enriched from Day5 to Day7 (Figure S3D). The regulons were then binarized to clarify interpretation. The pluripotent Epiblast-like population had largely lost primed pluripotency EPISC-specific regulons (e.g., Pou3f1) (Buecker et al., 2014) and reprogrammed with remarkably similar regulation to the ESC/ESC2CL with well-known naïve pluripotency regulons (e.g. Mybl2, Esrrb, Klf2, Klf4) and shared broader pluripotency continuum regulons (e.g., Sox2, Nanog, Pou5f1) with EPISC (Figure 3E) (Nichols and Smith, 2009; Weinberger et al., 2016). Not surprisingly, the ESC and ESC2CL populations had few differences at the regulatory level.

As anticipated, the PE-like and TE-like cells shared important extraembryonic regulons (e.g., Klf6, Kdm5a, Creb3, Elf1, Elf4) (Figure 3E) (Burton et al., 2013; Krendl et al., 2017; Rivron et al., 2018; Yang et al., 2013). Furthermore, the TE-like region had enriched significant TE-specific regulon activity (e.g, Gata2, Gata3, Ascl2, Pparg) (Figure 3E) including Cdx2 regulon (Figure S3D) (Home et al., 2017; Krendl et al., 2017; Ralston et al., 2010). Also, the PE-like region had some enriched PE-specific regulon activity including Gata6 and Hnf1b (Figure S3D) (Lo Nigro et al., 2017). Average gene expression for the same transcription factors of regulon analysis reflected a similar pattern (Figure 3F), although far less specific, highlighting the value of specific downstream regulation analyses when comparing cells (Aibar et al., 2017; Woogeng et al., 2020).

The gene regulation and RNA splice differences raised uncanny distinction of the regions. We investigated numerous mouse RNA Spliceosome genes (Kanehisa and Goto, 2000) and found some with discrete differences among the day-specific regions (Figure S4A). The Mbnl splice factors that repress naïve pluripotency-specific splicing (Han et al., 2013) were only active in the TE-like and PE-like region cells Figure S4A). Also, Mbnl3 is a core trophoblast gene induced by Gata3 and Cdx2 (Ralston et al., 2010), and was neatly expressed in the TE-like region. Interestingly, *Mbnl2* was one of the top20 markers for the TE-like region among other TE genes (Figure S4B).

Taken together, the three diverging regions of BCLH reprogramming cells formed over time with specific epigenetic splicing, expression, and downstream regulation that grossly reflected the three BC-cell lineages; such may explain the controlled order and development of BCLH observed in the cell culture plate (Kime et al., 2019).

### Some BCLH Cells Adopt the Regulatory Networks of Established Models

BCLH SES reprogramming induces many cells on the plate regulated spatially to resemble BCLHs (Kime et al., 2019). To better isolate BC-like cells *in silico*, we selected the induced cells that were most ICM/Epiblast-like (iEPI), TE-like (iTE), and PE-like (iPE) cells based on critical gene expression criteria (see methods). We also included comparable numbers of the ESCs/ESC2CL cells and starting EPISC. For established TE and PE model cell data, we sourced a loom file (Posfai et al., 2020) that was built from established reports’ scRNA-seq data. We then merged samples and normalized features and counts to integrate the data fairly (see methods).

Seurat UMAP clustering from gene expression generally showed distinct populations based on type although some batch effects were obviated separating TE samples that had different origins (Figure 4A). Some TE and PE cells clustered together, as was seen with some iTE and iPE cells, likely based on common extraembryonic expression. Excitingly, analysis with SCENIC showed distinct trends among putative similar cells (Figure S4C) and SCENIC binarized regulon analysis revealed, again, the distinction of three regions of BC-like lineage cells that resembled putative models but with more established BC-like cell factors (Figure 4B). The iEPI cells were regulated alike the ESC/ESC2CL, having lost nearly all EPISC-specific regulons and acquiring those of naïve pluripotency. The iPE were regulated to have PE regulons at similar or low levels and the iTE were regulated to have many of the TE regulons. Expectedly, the induced extraembryonic-like cells (iPE/iTE) had many regulons shared with the extraembryonic PE/TE cell models (Figure 4B) and some differences that appeared to come from batch effect. It was clear that the iPE and iTE samples had lost many EPISC regulons that were also not found in the PE and TE model data, although some remained (Figure 4B). Checking average gene expression for the transcription factors of the regulons expectedly showed a similar pattern with less distinction (Figure 4C), highlighting the importance and power of gene regulatory networks to determine a cell identity (Aibar et al., 2017). We imported the binarized regulon activity tables to the Seurat object and used FindMarkers to identify pluripotent, PE, and TE markers, select the top 10 of each, and plot the associated activity (Figure 4D). Indeed, nearly all top markers discovered were highly reported genes for their correlating cell states (e.g., Klf2, Mybl2, Nanog, Prdm14:: Cdx2, Ets2, Gata2, Gata3:: Sox17, Sox7, Gata4, Gata6), and were regulated relatively neatly among induced and model populations (Figure 4D). Notably, the iPE population had more iTE/TE extraembryonic regulons than the PE population (Figure 4B,D). In general, each of the three BC-like regions in BCLH appeared to be regulated by the critical transcription factors of their putative embryonic equivalents.

**Figure 4:**
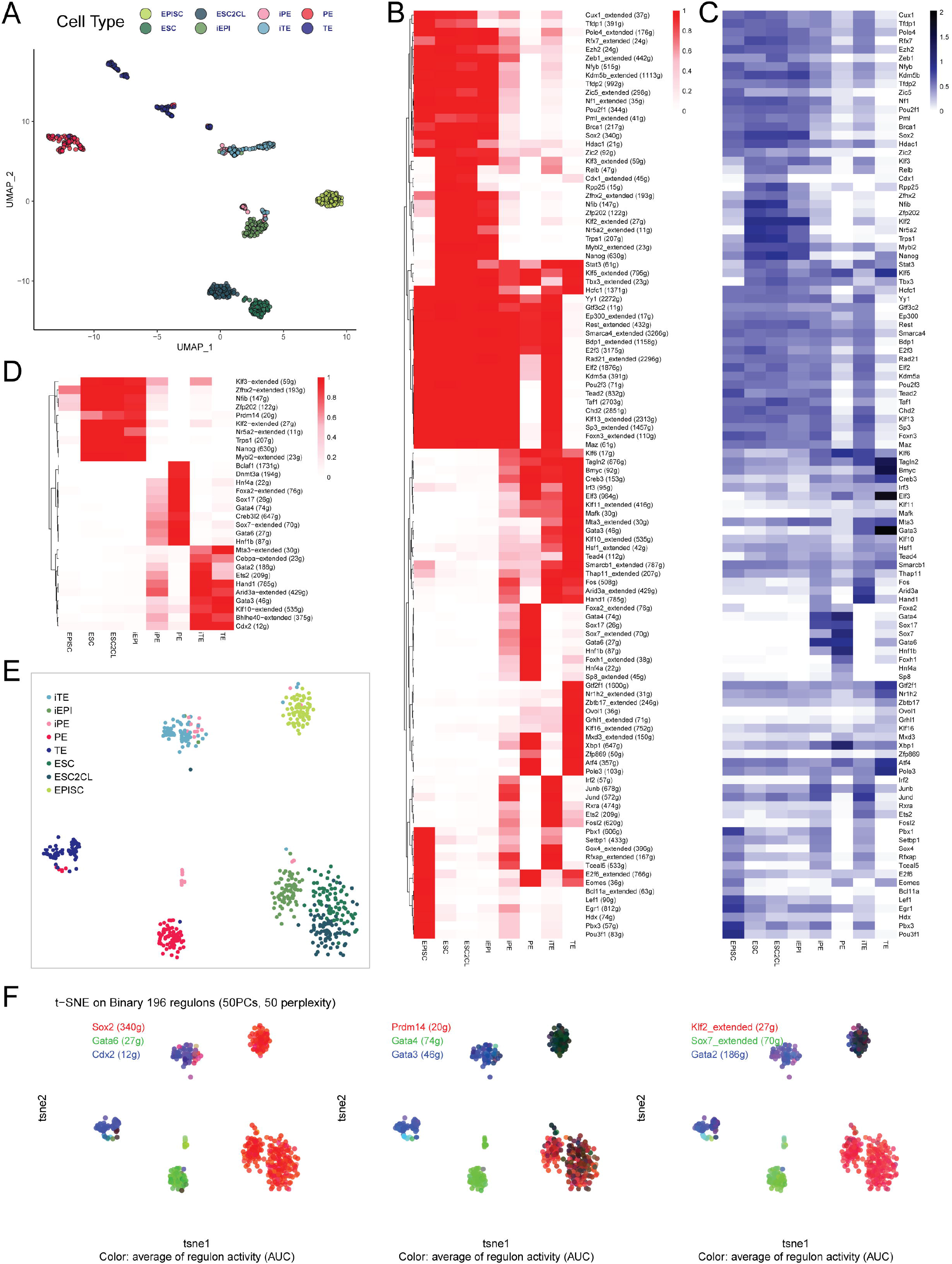
Select BCLH SES Cells Reprogram Meaningfully Close to Model Cells. **A)** UMAP based plot for EPISC, ESC, ESC2CL, iEPI, iPE, iTE, PE, and TE samples. **B)** SCENIC Binarized regulons with heatmap(pheatmap) regulon activity (red scale). **C)** Heatmap(pheatmap) of the row-matched transcription factor average expression (log-transformed) for the regulons of Figure 4B. **D)** Pluripotent, PE, and TE, combined Top 10 markers based on Binarized Regulons. **E)** SCENIC tSNE plot based on regulon activity for each sample. **F)** Average Regulon activity in RGB color for pluripotency regulons (red), TE regulons (green), and PE regulons (blue) across SCENIC tSNE plot (Figure 4E).

As shown previously, re-clustering cells based on SCENIC regulon activity can provide clearer identity-based clustering (Aibar et al., 2017; Posfai et al., 2020). Upon doing so in tSNE map, a pattern of cells reflecting the reprogramming and cell states had clarified (Figure 4E) and most of the batch effects between the TE cells was reduced, although some model PE and TE cells still mixed. As seen with the regulon heatmaps, the ESC and ESC2CL cells nearly shared the same tSNE space than when based on gene expression, strengthening the notion that these cells were more similar at the gene regulatory level when cultured in 2iLIF (Figure 4E); expectedly, the iEPI cells now clustered very close to the ESC/ESC2CL cells (Figure 4E). The iTE and iPE cells had dimensionally moved toward the TE and PE models with some iPE cells remarkably close to the PE model cluster (Figure 4E). To view individual BC-lineage critical regulon activity (Figure 4D), we prepared three tSNE plots with lineage-specific regulons on each plot that explained the regulatory activity responsible for the diverse cell types (Figure 4F).

At last, select cells of the three major regions of cells on the plate had iEPI, iPE, and iTE, cells present that had undergone remarkable cell reprogramming from primed pluripotent EPISC toward model cells of earlier embryos with sophisticated gene regulation at the regulon and RNA splicing levels.

## DISCUSSION

Until now we had seen self-assembly and order to resemble BCs in the BCLH SES (Kime et al., 2018, 2019). In addition to confirming those observations, this study provided numerous aspects of gene expression, RNA splicing regulation, and gene regulator network level that greatly strengthened our understanding that EPISC can reprogram to represent BCs. The numerous ZGA signature and germ program related genes provide further inspiration to wonder how the cell reprogramming is engaged although anticipated Dux and Zscan4 expression was not seen at these time points.

The BCLH SES is unique from its cell origin, defined conditions, and output. Given that the MERVL reporter activated early on, and Day5 cells neatly branched toward Day7, we anticipate that an earlier time-point of a unified reprogramming precursor cell may exist for exploit. Discovery and optimization of that cell, which reprograms from EPISC, may further advance the clarity and distinction of this SES. We anticipate that lineage tracing based on the Day5 data may help discern key precursors and emerging populations for study and optimization. We would also like to include early BC ICM cells and BC Epiblast-specific cells for scRNA-seq analysis that we could not presently access.

The MERVL reporter has had significant utility in our SESs and in the BCLH system it is more broadly induced than in the induced BC-like cysts (iBLC) (Kime et al., 2019) where the defined conditions are modified to different in phases. Conversely, BCLH SES cells proceed more rapidly to less organized BC-like hemispheres instead of puckering from the plate as floating self-organizing cysts. Since the MERVL reporter was highly active in all three regions of this study across Day5 and Day7, it provided some clues about unique early embryonic programs that may be engaging the genome. Surprisingly, the ESC2CL cells in our base condition with apparent *MERVL::RFP* expression had some interesting differences from ESCs in gene expression, but at the regulatory level they near-completely collapsed to the same cluster. We speculate that the MERVL reporter may have significant utility in cell reprogramming that had numerous ZGA related genes and we wonder if our ESC data challenges the reporter’s value in 2iLIF.

### Cell Reprogramming

In general, the reprogramming cells grossly lost their donor cell state and took on the programs of early BC-like cell lineages which reflected prior observations with high detail. The XGFP+ iEPI population was previously shown as readily potentiated for high contribution in chimeric embryos (Kime et al., 2016). Although the TE-like and PE-like regions could be identified and had convincing cells therein, we wonder if such cells could seed trophoblast stem cell or XEN cell cultures if transferred to appropriate conditions. In the BCLH SES, the TE-like region generally had low *Cdx2* expression despite the specific enrichment of the Cdx2 regulon, perhaps related to abundant keratin expression and TE-related transcription factor involvement. Interestingly, the TE can be specified independent of Cdx2 (Wu et al., 2010). Although Cdx2 is not required for blastocyst formation (Meissner and Jaenisch, 2006), its role in implantation is important and distinct CDX2+ TE is roundly regarded (Strumpf et al., 2005). The PE-like region was less distinguished and had significant overlap between the Epiblast-like and TE-like regions; perhaps the molecular distinction of the emerging iPE is as complicated as its natural ICM-to-extraembryonic transition. Traces of gene expression throughout this study suggested that iPE and PE-like region cells shared a pluripotent-like origin with the iEPI population reminiscent to the GATA4+, GATA6+, PDGFRA+ positive cells arising at the inner-face of BCLH pluripotent cells (Kime et al., 2019). Consideration for PE cells has weighed heavily on various SESs and we suspect that correct hypoblast formation will remain a hinge point for healthy embryoid development.

## AUTHOR CONTRIBUTIONS

Conceptualization, C.K.; Methodology, C.K., K.T.; Experimentation, C.K., H.H., H.S., Y.S.; Formal Analysis, C.K., K.T.; Investigation, C.K.; Resources, C.K., M.T., Y.S., K.T.; Writing – Original Draft, C.K.; Writing – Revision & Editing, C.K., K.T.; Visualization, C.K.; Project Supervision, C.K.; Bioinformatics: C.K.; Project Administration and Funding: C.K.

## ACKNOWLEDGEMENTS

We are grateful for the support from RIKEN Center for Integrative Medical Sciences for specific training and research environment with support from Erik Arner, Piero Carninci, Imad Abugessaisa, Akira Hasegawa, and Teruaki Kitakura. We also thank Osamu Nishimura and the Shigehiro Kuraku Lab at RIKEN for providing a HPC for Cell Ranger processing. We thank Eszter Posfai of Princeton University, and both Vincent Pasque and Adrian Janiszewski of Katholieke Universiteit (KU) Leuven for providing important control data and thoughtful discussions.

---

**Figure S1:**
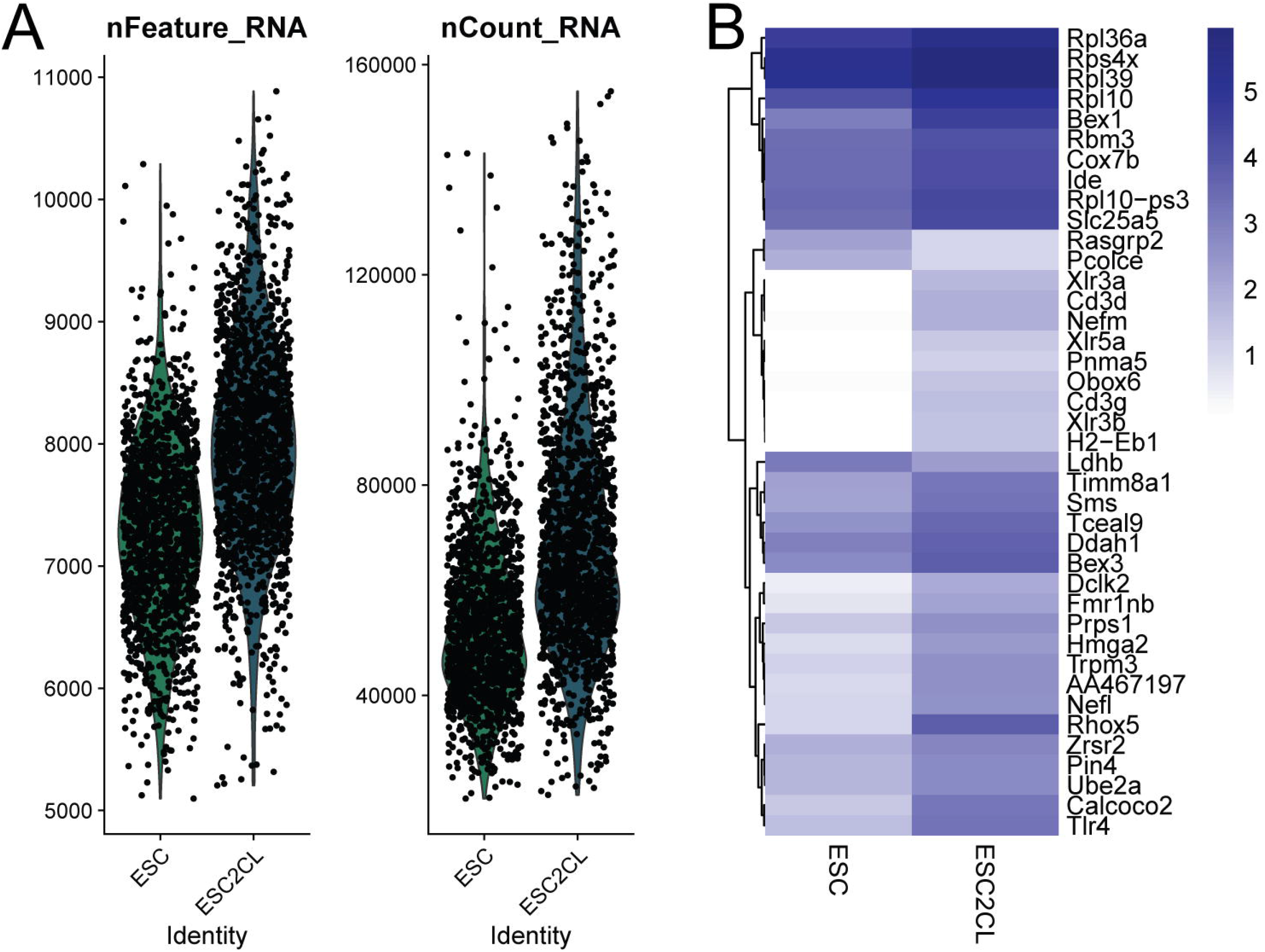
Related to Figure 1. **A)** Seurat violin plots for features and counts of ESC and ESC2CL samples. **B)** Heatmap(pheatmap) of the average gene expression of the ESC top 20 markers combined with ESC2CL top 20 markers.

**Figure S2:**
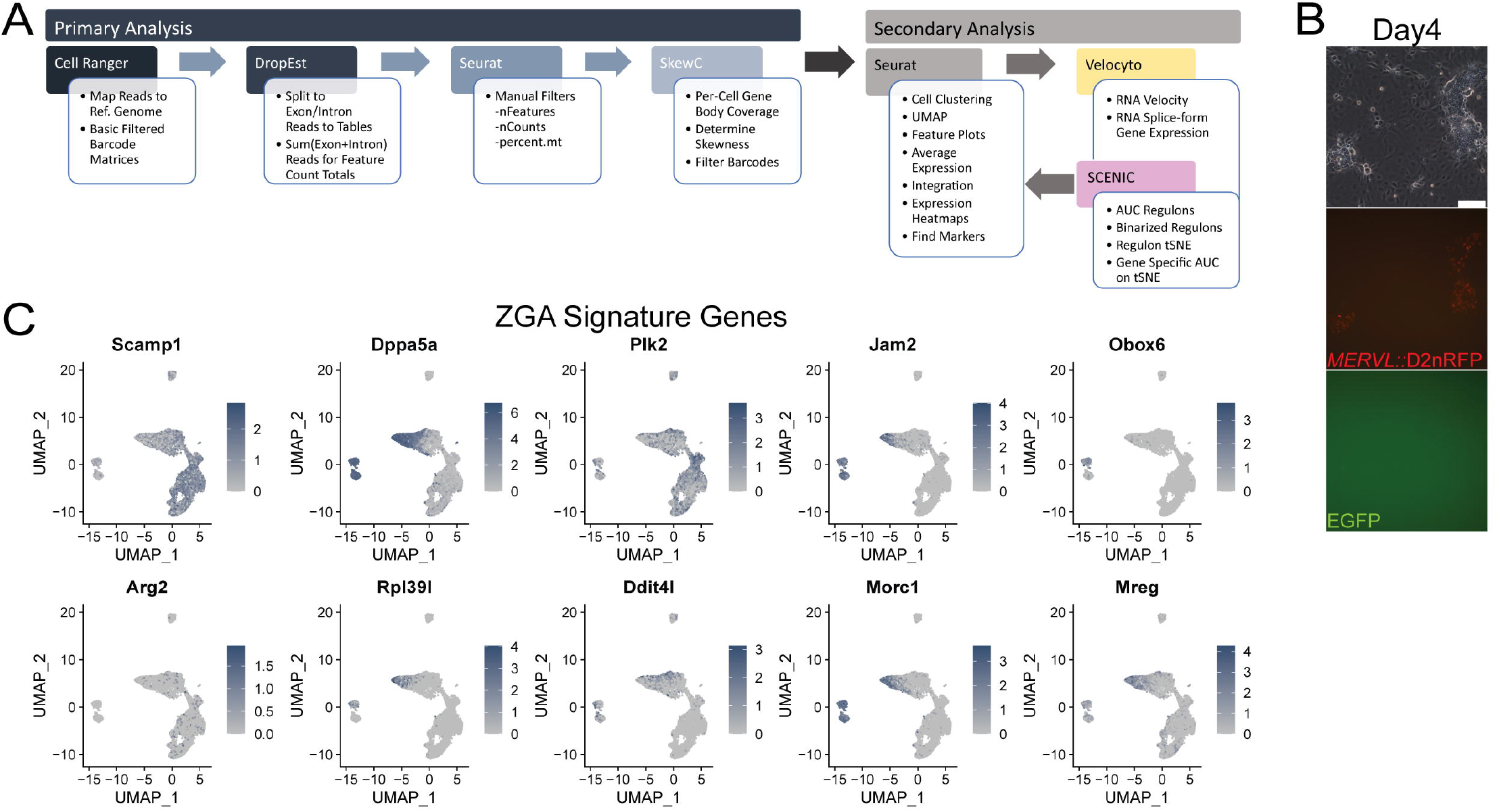
Related to Figure 2. **A)** An overview of the Primary and Secondary Analysis of scRNA samples in this study. **B)** BCLH reprogramming from EPISC with *MERVL::*D2nRFP and XGFP reporters imaged on Day4 for brightfield (top), *MERVL::*D2nRFP expression (mid), and XGFP expression (bot). *Scale bar = 100μm*. **K)** UMAP based gene expression feature plots for ZGA signature genes.

**Figure S3:**
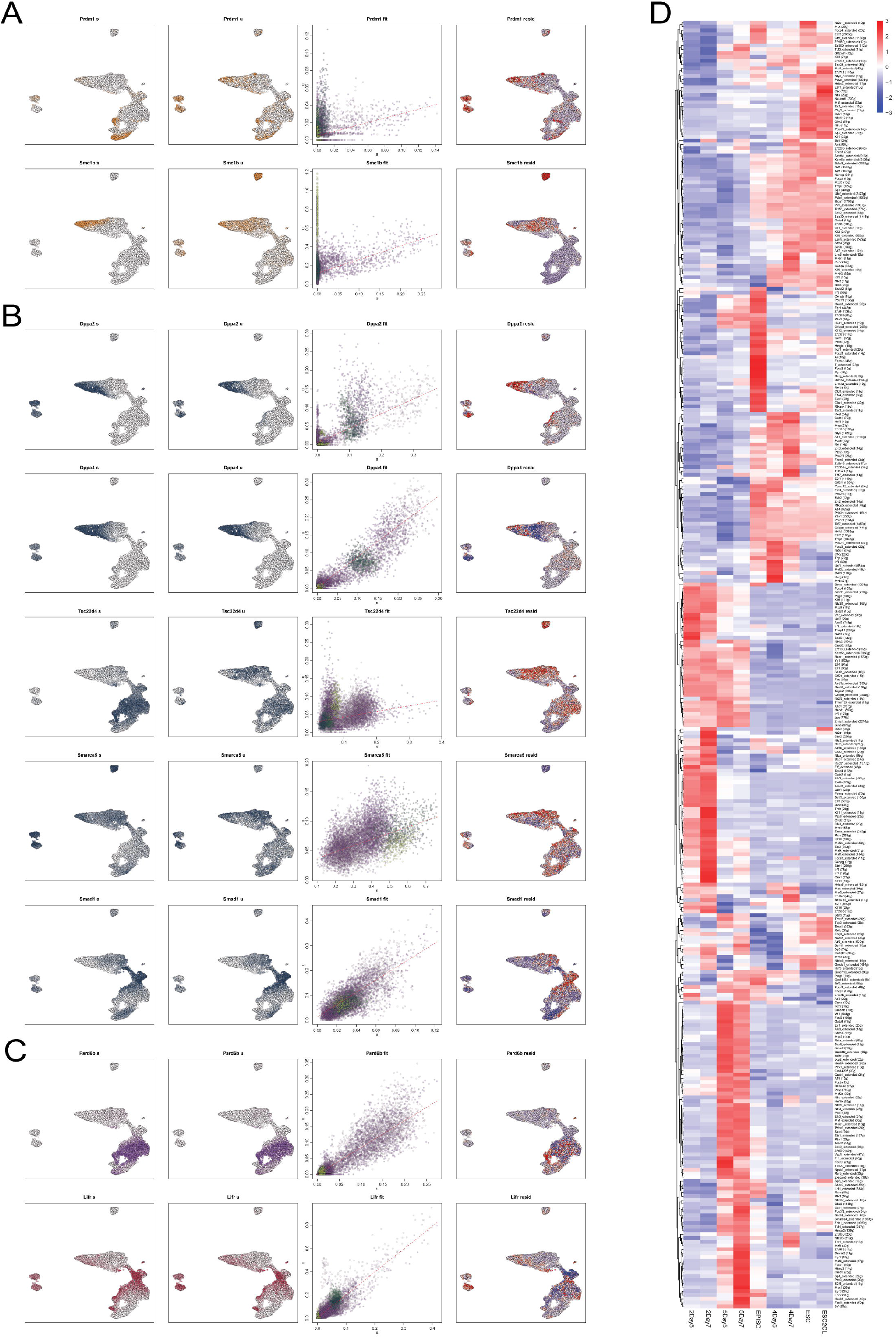
Related to Figure 3. **A)** UMAP based RNA splicing plots for spliced (s), unspliced (u) reads, with Cell Type coloring (Figure 3A) and residual (resid) unspliced expression shown for germ program factors. **B)** UMAP based RNA splicing plots for spliced (s), unspliced (u) reads, with Cell Type coloring (Figure 3A) and residual (resid) unspliced expression shown for germ ZGA-like regulators. **C)** UMAP based RNA splicing plots for spliced (s), unspliced (u) reads, with Cell Type coloring (Figure 3A) and residual (resid) unspliced expression for TE (purple) and PE (burgundy) genes. **D)** SCENIC total AUC regulon activity by day-specific cells and control cells.

**Figure S4:**
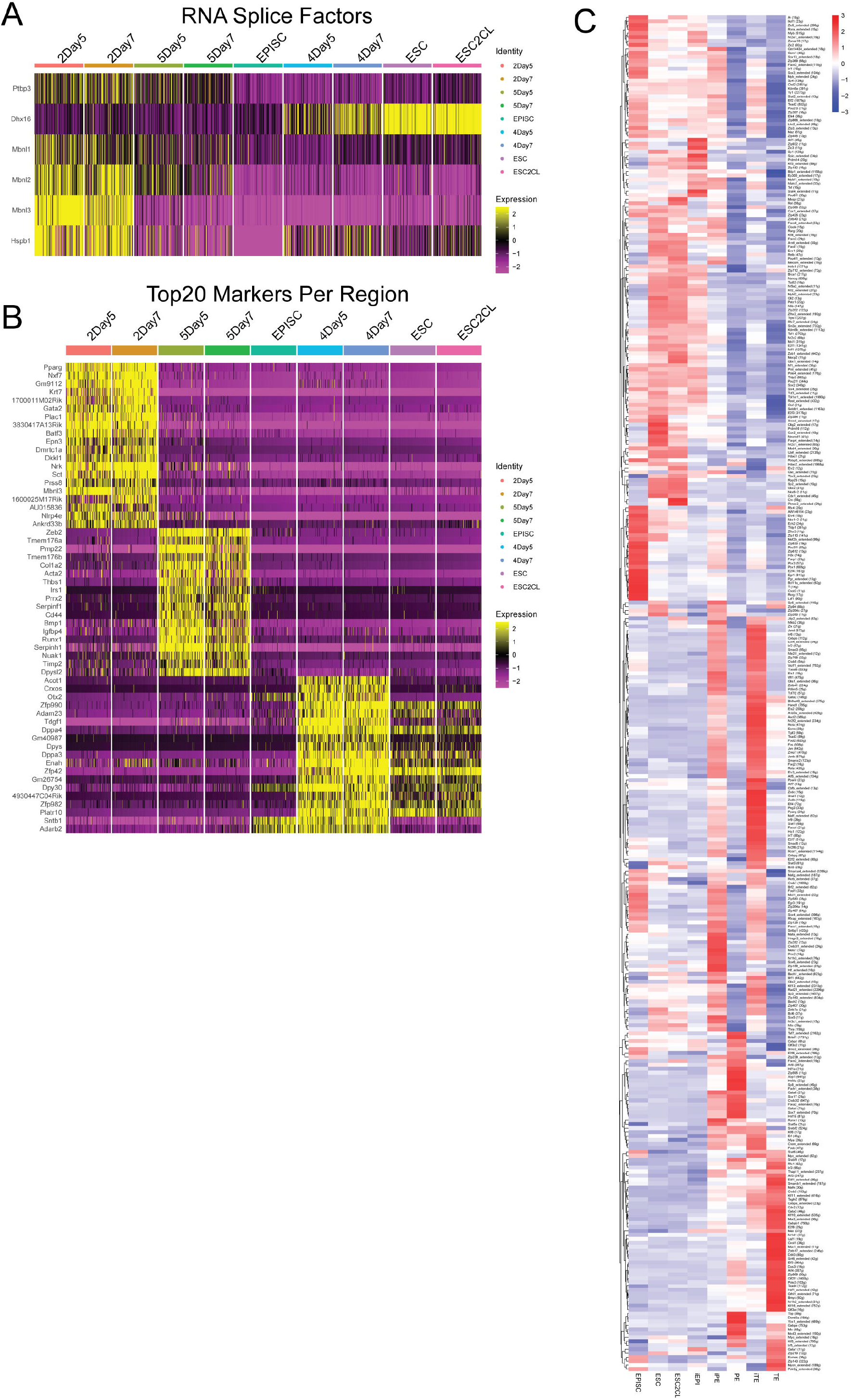
Related to Figure 3, Figure 4. **A)** Heatmap(DoHeatmap) plot of notable RNA splicing (spliceosome) factors of all cells, ordered by the day-specific cells and control cells. **B)** Heatmap(DoHeatmap) plot of the combined top20 markers per region (2: TE-like, 5: PE-like, and 4: EPI-like) ordered by day-specific cells and control cells. **C)** SCENIC total AUC regulon activity for select iTE, iPE, and iEPI, and control TE, PE, ESC/ESC2CL, and EPISC.

**Table S1: ESC Markers**

ESC markers discovered from FindMarkers in Seurat.

**Table S2: ESC2CL Markers**

ESC2CL markers discovered from FindMarkers in Seurat.

## EXPERIMENTAL PROCEDURES

### EPISC Culture and BCLH Reprogramming

EPISCs culture and reprogramming was performed as described previously (Kime et al., 2016, 2019), and reprogramming included Sodium Pyruvate.

### ESC Culture and MERVL Reporter Integration

BL6 ESCs were converted from 3iLIF conditions and cultured in 2iLIF conditions with CTSFES basal media as described previously (Kime et al., 2019) on iMatrix511 coated 6-well plates. Cells were integrated and selected for *MERVL::RFP* reporters (mCherry and D2nmCherry) in piggyback vectors the same as with EPISC in our previous study (Kime et al., 2019). After several passages, the two different populations of cells became obvious and stabilized.

### scRNA-seq Sampling and Processing

Cells were dissociated and passed through cell screen cuvettes to isolate mostly healthy single-cells that were prepared with 10x Chromium Single Cell 3’ Library & Gel Bead Kit V3.0. Sample libraries were finalized and sequenced on one Hiseq X lane (150bp PE; Macrogen) for each. Standard Cell Ranger protocol detected sample chemistry and produced ‘possorted’ BAM files from which the subsequent Primary Analysis workflow in Figure S2A was performed.

To match loom data counts for TE and PE control cells (Posfai et al., 2020), we prepared our data tables similarly with DESeq2 size factor normalization prior to merging samples. The iEPI, iTE, and iPE cells were selected with the following criteria:

iEPI: Zfp42 > 0.1 & Klf2 > 0.1 & EGFP > 0 & Prdm14 > 0.001
iTE: Cdx2 > 0 & Gata2 > 0
iPE: Gata6 > 0 & (Sox7 > 0 | Gata4 > 0 | Sox17 > 0)

### Microscopy

Brightfield and live cell RFP and GFP fluorescence was imaged with a Olympus IX71 Microscope.

